# Reciprocal changes in voltage-gated potassium and subthreshold inward currents help maintain firing dynamics of AVPV kisspeptin neurons during the estrous cycle

**DOI:** 10.1101/2021.07.14.452390

**Authors:** J. Rudolph Starrett, R. Anthony DeFazio, Suzanne M. Moenter

## Abstract

Kisspeptin-expressing neurons in the anteroventral-periventricular nucleus (AVPV) are part of a neural circuit generating the gonadotropin-releasing hormone (GnRH) surge. This process is estradiol-dependent and occurs on the afternoon of proestrus in female mice. On proestrus, AVPV kisspeptin neurons express more kisspeptin and exhibit higher frequency action potentials and burst firing compared to diestrus, which is characterized by a pulsatile rather than a prolonged surge of GnRH secretion. We hypothesized changes in voltage-gated potassium conductances shape activity profiles of these cells in a cycle-dependent manner. Whole-cell voltage-clamp recordings of GFP-identified AVPV kisspeptin neurons in brain slices from diestrous and proestrous mice revealed three subcomponents of the voltage-sensitive K^+^ current: fast-inactivating, slow-inactivating, and residual. During proestrus, the V50 of inactivation of the fast-inactivating current was depolarized and the amplitude of the slow-inactivating component was reduced compared to diestrus; the residual component was consistent across both stages. Computational models were fit to experimental data, including published estrous-cycle effects on other voltage-gated currents. Computer simulations suggest proestrus-typical K^+^ currents are suppressive compared to diestrus. Interestingly, larger T-type, persistent-sodium, and hyperpolarization-activated currents during proestrus compensate for this suppressive effect while also enabling post-inhibitory rebound bursting. These findings suggest modulation of voltage-gated K^+^ and multiple subthreshold depolarizing currents across the negative to positive feedback transition maintain AVPV kisspeptin neuron excitability in response to depolarizing stimuli. These changes also enable firing in response to hyperpolarization, providing a net increase in neuronal excitability, which may contribute to activation of this population leading up to the preovulatory GnRH surge.

**Significance Statement:** GnRH neurons provide the central signal to initiate ovulation by releasing a surge of hormone. GnRH neurons are regulated by other cells including those expressing kisspeptin, a potent stimulator of GnRH secretion. Kisspeptin neurons in the anteroventral-periventricular nucleus (AVPV) express more kisspeptin and become more active during the afternoon of proestrus, the phase of the rodent estrous (reproductive) cycle when the GnRH surge occurs. We found voltage-dependent potassium currents in AVPV kisspeptin neurons change with phase of the estrous cycle. Firing simulations indicated these changes are suppressive if occurring in isolation. But proestrous-typical increases in subthreshold depolarizing currents overcome this suppression and promote greater excitability by increasing rebound firing, possibly contributing to the preovulatory activation of this system.

## Introduction

The anteroventral-periventricular (AVPV) nucleus is a critical site involved in the regulation of female fertility, specifically the control of the estradiol-dependent process of ovulation (Kalra and McCann, 1975; Goodman, 1978; Wiegand and Terasawa, 1982; Petersen and Barraclough, 1989; Petersen et al., 1989). The AVPV contains neurons that express both estrogen receptor α (ERα) and kisspeptin (Simerly, 1998; Smith et al., 2005), a neuropeptide that acts at gonadotropin-releasing hormone (GnRH) neurons to potently induce GnRH neuron activity and hormone secretion (Han et al., 2005; Messager et al., 2005; Pielecka-Fortuna et al., 2008; Zhang et al., 2008; Caraty et al., 2013). Ovulation is initiated by positive feedback actions of estradiol at the central and pituitary levels (Döcke and Dörner, 1965; Sarkar et al., 1976; Moenter et al., 1991). In a prevailing theory, sustained elevation of circulating estradiol during the late follicular phase (the cycle day of proestrus in rodents) exerts positive feedback effects via ERα that drive increased kisspeptin expression in AVPV neurons (Smith et al., 2005; Gottsch et al., 2006; Oakley et al., 2009). Action potential firing by AVPV kisspeptin neurons is also increased on proestrus vs diestrus (a time of homeostatic negative feedback) (Wang et al., 2016). Combined, these effects are thought to increase kisspeptin secretion onto the GnRH neuron, as increases in firing frequency are typically associated with increased neuropeptide release (Dutton and Dyball, 1979; Cropper et al., 2018). Estradiol positive feedback is likely conveyed to GnRH neurons by increased kisspeptin signaling, causing a surge of GnRH release which stimulates a luteinizing hormone (LH) surge from the pituitary. The LH surge stimulates ovulation (Greep et al., 1942).

If the above theory is correct, the firing activity of AVPV kisspeptin neurons is a critical piece of this surge induction process, serving as a gating mechanism controlling kisspeptin release and thus the physiologic cascade culminating in ovulation. Several studies have focused upon the electrical properties of these cells and how they change across the reproductive cycle, measured via recordings performed in acutely prepared mouse brain slices during periods of estradiol negative (typically diestrus) vs. positive (proestrus) feedback. AVPV kisspeptin neurons recorded on proestrus maintain higher firing rates than on diestrus even when fast synaptic transmission was blocked, suggesting a shift in intrinsic properties across the diestrus-proestrus transition enables increased firing during proestrus (Wang et al., 2016). Voltage-gated conductances may provide a basis for these shifts in activity since they help control membrane potential and shape firing dynamics in electrically-excitable cells. T-type calcium, persistent sodium, and hyperpolarization-activated currents were all found to be larger in AVPV kisspeptin neurons during proestrus (Piet et al., 2013; Zhang et al., 2015; Wang et al., 2016). Increases in these depolarizing currents, which can be active at subthreshold membrane potentials, could promote greater firing activity. Potassium currents, which strongly influence firing onset of many neurons and are vital for proper repolarization of the membrane (Hille, 2001), remained uncharacterized in AVPV kisspeptin neurons. K^+^ currents change across the estrous cycle and in response to estradiol in GnRH neurons and kisspeptin neurons in the arcuate nucleus (DeFazio et al., 2002, 2019; Pielecka-Fortuna et al., 2011).

We hypothesized voltage-gated potassium currents in AVPV kisspeptin neurons are regulated by estrous cycle stage. Classic voltage-clamp approaches revealed three conductances contribute to the total voltage-gated potassium current in these cells, and each component was characterized during the afternoon of diestrus (negative feedback) and proestrus (positive feedback). It can be difficult to predict how multiple voltage-gated conductances interact to control the membrane voltage. Computational modeling studies were thus conducted to better understand how estrous cycle modulation of these and other currents influence action potential firing output.

## Materials and Methods

### Animals

The University of Michigan Institutional Animal Care and Use Committee approved all procedures. Adult female Kiss1-hrGFP mice (Cravo et al., 2011) which express humanized *Renilla* GFP under control of the kisspeptin promoter, were used for these studies. Mice were provided with Harlan 2916 chow and water *ad libitum* and were held on a 14L:10D light cycle with lights on at 3:00 AM Eastern Standard Time. Estrous cycle stage was monitored by vaginal cytology for at least one week before experiments. Uterine mass was measured after brain slice preparation to confirm cycle stage. Uterine mass >100mg indicated *in vivo* exposure to a high concentration of estradiol, typical of proestrus, whereas mass <60mg indicated exposure to low estradiol, typical of diestrus (Shim et al., 2000).

### Experimental Design

Brain slices were prepared from cycling adult female mice (age 55-154 days) during cycle stages corresponding to estradiol negative feedback (afternoon of diestrus) or positive feedback (afternoon of proestrus). Whole-cell voltage-clamp recordings of hrGFP-identified AVPV kisspeptin neurons were used to characterize macroscopic voltage-gated potassium currents; three currents (fast transient, slow transient, residual) were identified and separated using pharmacological and/or voltage-based subtraction methods for characterization. Current-clamp recordings were used to monitor action potential firing. Computational modeling was used to predict how each type of potassium current regulates AVPV neuron excitability, and how changes in multiple voltage-gated currents across the estrous cycle may impact excitability.

### Brain slice preparation

Chemicals were purchased from Sigma-Aldrich unless noted. All solutions were bubbled with 95% O2/5% CO2 for at least 15 minutes before exposure to tissue. All mice were euthanized at 3:00 to 4:00 PM Eastern Standard Time, and the brain was rapidly removed and placed in ice-cold sucrose saline solution containing the following (in mM): 250 sucrose, 3.5 KCl, 25 NaHCO_3_, 10 D-glucose, 1.25 Na_2_HPO_4_, 1.2 MgSO_4_, and 3.8 MgCl_2_, at pH

7.6 and 345 mOsm. Coronal (300 µm) slices were cut with a VT1200S Microtome (Leica Biosystems). Slices were incubated in a 1:1 mixture of sucrose saline and artificial cerebrospinal fluid (ACSF) containing (in mM) 135 NaCl, 3.5 KCl, 26 NaHCO_3_, 10 D-glucose, 1.25 Na_2_HPO_4_, 1.2 MgSO_4_, and 2.5 CaCl_2_ at pH 7.4 and 305 mOsm for 30 min at room temperature (∼21 to 23 °C) and then were transferred to 100% ACSF for an additional 30-180 min at room temperature before recording. For recording, slices were placed into a chamber and perfused (3 ml/min) with carboxygenated ACSF kept at 31°C with an inline heating unit (Warner Instruments). GFP-positive AVPV kisspeptin neurons were identified by brief illumination at 488 nm on an Olympus BX51WI microscope. Recordings were performed 1-4 h after brain slice preparation. No more than four cells were recorded per mouse; data values from cells from the same animal were not clustered in a manner that would typically contribute to reduced variability.

### Voltage-clamp recordings

Recording micropipettes were pulled from borosilicate capillary glass using a Flaming/Brown P-97 puller (Sutter Instruments) to obtain pipettes with a resistance of 1.5-3.5 MΩ when filled with pipette solution, which consisted of (in mM) 135 K-gluconate, 10 KCl, 10 HEPES, 5 EGTA, 0.1 CaCl_2_, 4 MgATP, and 0.4 NaGTP, 305 mOsm and pH 7.2 with NaOH. Pipettes were wrapped in Parafilm to reduce capacitive transients. All potentials reported were corrected on-line for a liquid junction potential of -15.7 mV(Barry, 1994). Recordings were performed with one channel of an EPC-10 dual patch-clamp amplifier and PatchMaster software (HEKA Elektronik). After achieving the on-cell configuration with seal resistance >2.0 GΩ, fast capacitive transients were minimized and the whole-cell configuration was achieved by rupturing the cell membrane with brief suction. The membrane potential was held at -70 mV between voltage-clamp protocols. Passive membrane properties and voltage-clamp quality were calculated from the averaged current response (after on-cell capacitive current subtraction) to sixteen -5 mV, 20 ms test pulses from a holding potential of -70 mV performed immediately before and after each protocol. To ensure adequate recording quality, the following criteria were required for inclusion for analysis: uncompensated series resistance (Rs) <20 MΩ, input resistance (Rinput) >500 MΩ, capacitance (Cm) between 8 pF and 30 pF, holding current (Ihold) between -60 and 10 pA. Rs was compensated 50-85% for all protocols and recordings were excluded from analysis if >20% change in Rs occurred during the experiment.

To isolate voltage-gated potassium currents, TTX (2.5 µM, Tocris Bioscience), CdCl_2_ (100 µM), and NiCl_2_ (300 µM) were included in the ACSF to block Na^+^ and Ca^2+^ currents. Picrotoxin (100 µM) was used to block GABA_A_-receptor-mediated currents and D-2-amino-5-phosphonovalerate (APV; 20 µMm, Tocris Bioscience), and 6-cyano-7-nitroquinoxaline-2,3-dione (CNQX; 10 µM) were used to block ionotropic glutamate receptor-mediated currents.

### Total K^+^ current

In initial recordings of the total voltage-gated potassium current, two temporally-distinct peaks were visible in voltage steps from -100 to -10 or 0 mV. Further, a persistent negative slope in the outward current suggested a component with slow inactivation was present at potentials depolarized to -10 mV. Lengthening the voltage-clamp protocols allowed for more complete removal or induction of inactivation before the test pulse. To measure the voltage dependence of activation of the total voltage-gated K^+^ current, the membrane potential was held at -100 mV for 5 s to remove inactivation then stepped to test pulses of -70 to +40 mV (10 mV increments) for 1 s. To measure the voltage-dependence of inactivation, the membrane was held at potentials ranging from -100 to -10 mV (10mV increments) for 10s and then stepped to a test pulse of 0 mV for 1 s. The long duration of these protocols makes P/N subtraction (Armstrong and Bezanilla, 1977) unfeasible. Instead, slow capacitance and leak current subtraction were performed offline by subtracting the appropriately scaled average current response to one hundred -5 mV test pulses with capacitance and series resistance compensation activated (Kimm et al., 2015). Averages were scaled for each voltage step used in the protocol.

### Slow-transient K^+^ current

To characterize the slow transient K^+^ current, the same activation/inactivation protocols as above were repeated with 5 mM 4-aminopyridine (4-AP) included in the ACSF to reduce primarily the fast-transient current. To measure the time course of inactivation of the slow-transient K^+^ current, the membrane was held at -100 mV for 5 s to remove inactivation, then at -10 mV for durations varying from 0-8 s, followed by a test pulse at 0 mV for 1 s. To measure the time-dependence of recovery from inactivation, the membrane was held at -10 mV for 10s to inactivate the current, then at -100 mV for durations varying from 0-8 s, followed by a test pulse at 0 mV for 1 s. Offline leak subtraction was used as for recordings of the total K^+^ current (Kimm et al., 2015).

### Fast-transient K^+^ current

The fast-transient K^+^ current was isolated and quantified using a voltage-based subtraction method after reducing primarily the slow-inactivating and residual components with ACSF in which 20 mM tetraethylammonium chloride (TEA) replaced 20 mM NaCl. To measure the voltage-dependence of activation, two protocols (A and B) were run in series. For protocol A, the membrane potential was held at -100 mV for 200 ms, then current was measured during 150 ms test pulses from -70 to +40 mV (10 mV increments). Protocol B was identical to protocol A, except that the membrane potential was held at -30 mV rather than - 100 mV during the first 200 ms of the protocol. This fully inactivates the fast-transient current, isolating any remaining residual component during the test pulse. The current response during the test pulse of protocol B was subtracted from that of protocol A to yield the fast-transient current. To measure the voltage-dependence of inactivation, the membrane was held at -100 to -20 mV (10 mV increments) for 200 ms, before stepping to a test pulse at -10 mV (150 ms). Current occurring after a step from -30 mV holding potential was defined as residual current and was subtracted from earlier sweeps to yield isolated fast transient current. To measure the rate of inactivation of the fast-transient K^+^ current, inactivation was fully removed by holding the membrane at -100 mV for 300 ms. The membrane was then stepped to -30 mV for 0-250 ms followed by a test pulse of 0 mV for 100 ms. The component of the current that did not inactivate (residual current) was subtracted from the prior series to isolate the fast-transient current. To measure the rate of the recovery from inactivation, the fast-transient current was fully inactivated by holding the membrane at 0 mV for 300 ms, then recovery prepulses at -100 mV were applied for 0-250 ms before measuring current at 0 mV (100 ms). The current after the 0 ms recovery prepulse was subtracted from each test pulse to isolate the fast-transient K^+^ current. Slow capacitance and leak currents were corrected using P/-5 online leak subtraction with Vhold = -70 mV (Armstrong and Bezanilla, 1977).

It was not possible to quantify all parameters from every cell because not all cells remained within our quality control standards long enough for inclusion in the analysis of the time dependence of activation and inactivation. The number of cells and animals per measurement are shown in Tables 2 and 3

Total, fast-transient, slow-transient and residual K^+^ current densities were calculated by dividing by capacitance. Peak values were divided by the driving force according to the Goldman-Hodgkin-Katz (GHK) current equation using the calculated reversal potential (−94 mV) (Clay, 2000, 2009). Conductance values were then normalized to the maximum conductance (g_max_) to generate steady state activation/inactivation curves, which were fit with a Boltzmann function to calculate voltage of half (V50) activation or inactivation, and steepness (k). Inactivation and recovery from inactivation curves were fit to single exponentials to measure time constants *(τ)*.

### Current-clamp recordings

Depolarizing (2 to 28 pA, 2 pA increments) and hyperpolarizing (−25 to -5 pA, 5 pA increments) current pulses (500 ms) were applied in whole-cell configuration. The maximum amplitude of hyperpolarizing stimuli was limited to avoid hyperpolarizing the membrane beyond -100 mV, as further hyperpolarization was detrimental to recording stability. The targeted initial membrane potential was within 2 mV of -70 mV, which is close to the baseline membrane potential of these cells (DeFazio et al., 2014). Spikes were counted as action potentials only if their peak value was depolarized relative to -10 mV; spike number was plotted as a function of current injection.

### Modeling

#### Physiological data utilized

Mean ± SEM normalized activation, inactivation, recovery, and rate of inactivation curves were calculated for the total, fast-transient (fast), slow-transient (slow) and residual (resid) K^+^ current densities in each group. Leak-subtracted current traces from activation protocols were averaged for each current at each voltage step to obtain mean ± SEM current traces used for fitting. T-type calcium and persistent sodium conductance data were collected from published measurements, which indicated larger amplitudes during proestrus (Zhang et al., 2015; Wang et al., 2016). The hyperpolarization-activated conductance was estimated based on the amplitude and time course of the sag potential, which has larger amplitude during proestrus (Piet et al., 2013; Wang et al., 2016). Neurons were modeled as a single compartment with whole-cell capacitance of 17.62 pF based on the mean of experimentally-observed values (Table 1).

**Table 1.**
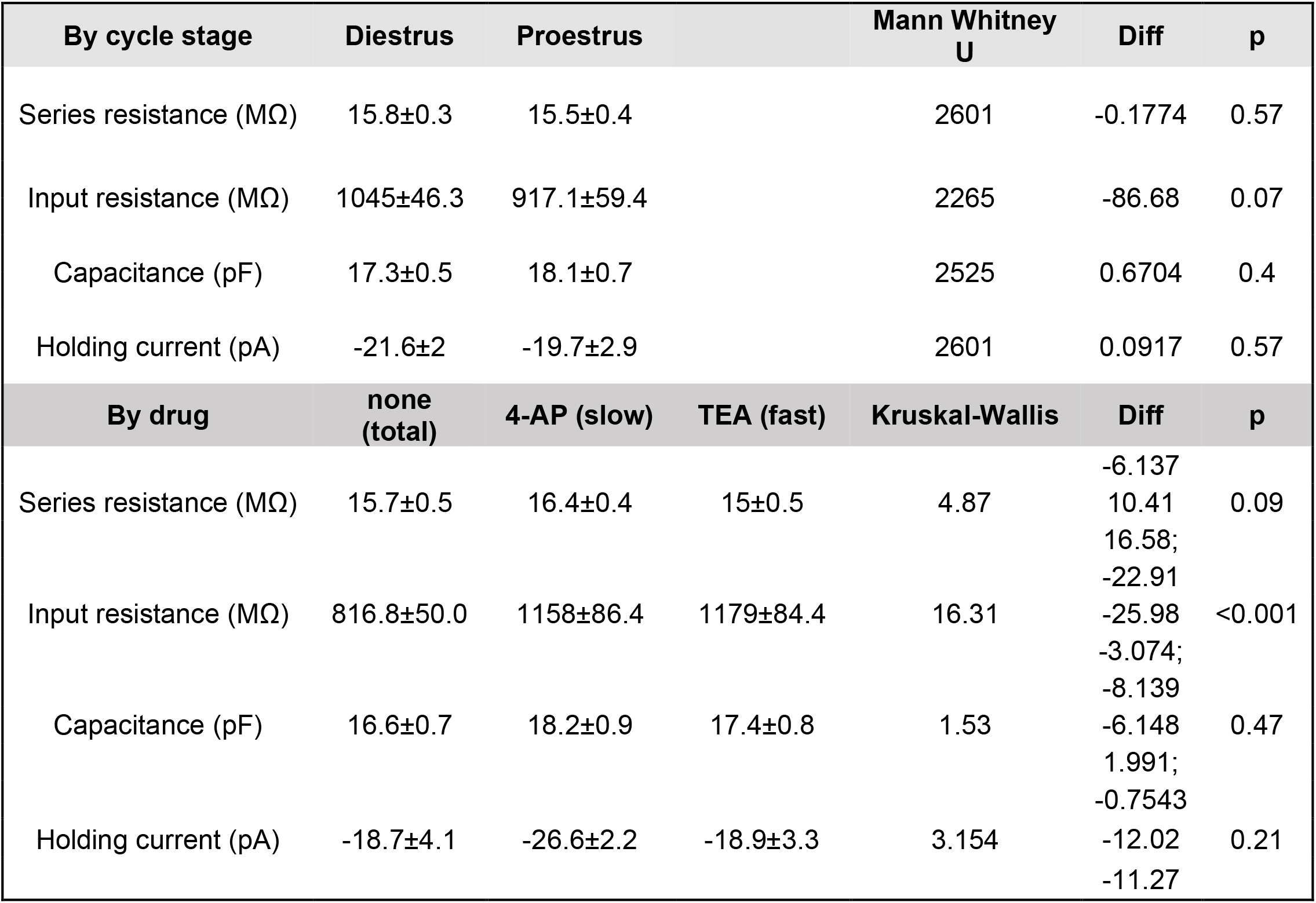
Passive properties and statistical comparisons

#### Equations

Na^+^, Ca^2+,^ and hyperpolarization-activated currents were modeled based on Ohmic driving forces (Eq 1-2). K^+^ currents were modeled based on nonlinear driving forces described by the Goldman-Hodgkin-Katz (GHK) current equation (Clay, 2000, 2009) (Eq 3).

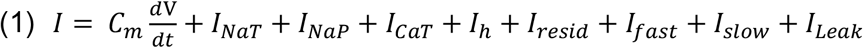

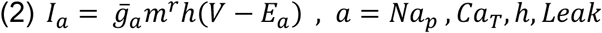

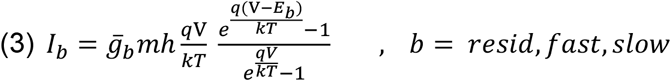

I is current (nA) across the neuronal membrane, C_m_ is the cell capacitance (pF), *V*_*m*_ is the membrane potential (mV), *t* is time (ms),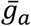 is the specific conductance (nS), *E*_*a*_ is the reversal potential (mV) of a given ion, *q* is the elementary charge, *k* is the Boltzmann constant, *T* is the absolute temperature, and *m* and *h* represent activation and inactivation variables, respectively. Exponents *r* represents the number of activation particles. These variables are governed by the following differential equations (Eq 4,5)

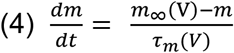

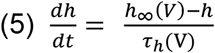

where *m*_*∞*_*(V)* and *h*_*∞*_*(V)* are the steady-state activation/inactivation functions for each variable and *τ*_*m*_ and *τ*_*h*_ are functions determining the voltage-dependent time constants (in ms) of activation/deactivation and inactivation/recovery, respectively. These functions take the form (Eq 6-9). *A*-*G* are free parameters adjusted during fitting.

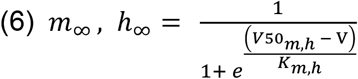

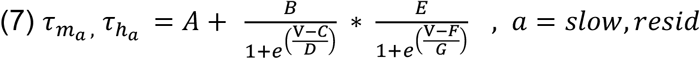

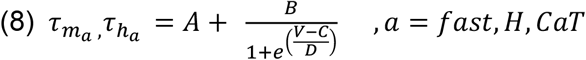

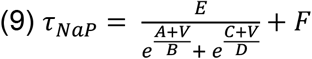

The transient sodium current (*Na*_*T*_) was modeled used a Markov chain with three states, open (*O*), closed (*C*), or inactivated (*I*). Transitions between states were either voltage sensitive or constants. The model used was modified from Adams et al., 2018. (Eq. 10-14).

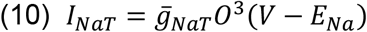

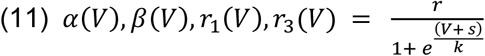

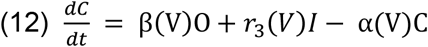

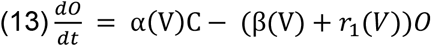

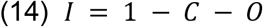

#### Parameter optimization

Simulated voltage-clamp experiments were performed using Xolotl (Gorur-Shandilya et al., 2018). Voltage-step protocols matched those used experimentally for each conductance. Current traces generated by models were analyzed in the same manner as the slice recording data. For each voltage step, peak currents were converted to conductance and normalized to generate activation/inactivation curves. For K^+^ models, permeability as calculated by the GHK current equation was used (Clay, 2000, 2009). *m*_*∞*_ and *h*_*∞*_ function parameters were adjusted manually until the normalized activation/inactivation curves were well fit to the experimental data. τ_*m*_, τ_*h*_, and 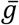 parameters were also adjusted so that the rise and decay (where applicable) phases of the current trace and peak amplitudes were well fit to the data. This process was iterative and was greatly facilitated by xolotl’s “ puppeteer” function, which allowed for rapid-recalculation and visualization of simulation results following parameter adjustment. Separate models were generated for each experimental group to reproduce estrous cycle effects on each of the voltage-gated currents.

The total K^+^ current was fit by simulating voltage-clamp of a cell containing the three K^+^ conductance subcomponents (fast, slow, resid) as they were fit to their respective drug-separated recordings. To better fit the total K^+^ recording,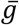 parameters were optimized using particle swarm optimization (Global Optimization Toolbox, Mathworks), which sought to minimize the sum of squares error between the current trace generated by the model and the mean data for the respective conductance.

#### Simulation of excitability

To study action potential firing, diestrous and proestrous neuron models were built with an identical sodium channel model. This model was made the same between groups because although there are no experimental data characterizing this conductance in AVPV kisspeptin neurons, action potential threshold (defined as when rate of rise exceeds 2V/s) and amplitude were the same between cycle stages (Wang et al., 2016), and these characteristics are highly influenced by transient sodium current properties (Kress et al., 2010).

To examine how conductances alter firing, a suite of conductance models corresponding to the respective experimental group were added to single-compartment neuron models, and the voltage of the model was allowed to vary, i.e., “ current clamp” . Using the same protocols from *in vitro* recordings, firing was stimulated by applying increasing current steps (−30 to 0 pA at 10 pA intervals, 2-28 pA at 2 pA intervals) to the model cell at a baseline membrane potential of -70 mV. The number of spikes fired during the stimulation is plotted against the stimulation amplitude to generate a firing-current (F-I) curve. To achieve this baseline potential, holding current and leak conductance were adjusted so that the membrane potential prior to the start of the current pulse was within 50 µV of -70 mV and input resistance was 1 GΩ (passive properties in Table 1). For diestrous and proestrous models, the simulated rheobase action potential waveform was compared to the peak-aligned mean rheobase action potential of the corresponding experimental group. This fit was improved by manually adjusting sodium conductance model parameters; adjustments were kept the same for the diestrous and proestrous models. To test predictions regarding estrous cycle effects or the role and/or interaction of various conductances, individual or multiple conductances fit to diestrous data were substituted into the proestrous model (and *vice versa*) for a particular simulation.

#### Statistics

Data were analyzed using Prism 9 (GraphPad). Shapiro-Wilk was used to test distribution of data. Parameters (mean ± SEM) were compared between cycle stages with unpaired statistical tests appropriate for experimental design and data distribution as indicated in the results section. The number of cells per group is indicated by n. For two-way ANOVAs and Mann-Whitney, difference of the means (Diff) was defined for cycle stage (diestrus– proestrus). For Kruskal-Wallis tests (Table 1), mean rank differences are reported as (total – slow), (total-fast), (slow – fast). Significance was set at *p* < 0.05.

## Results

### Passive properties

Series resistance, input resistance, capacitance, and holding current measurements were made before and after each protocol and values were averaged for each cell. There were no statistical differences between cells recorded during diestrus vs proestrus for any of the measured properties (Table 1). Comparing across drug treatments that are discussed below, input resistance was higher during 4-AP or TEA treatment (Table 1). Dunn’s multiple comparisons *post hoc* tests of input resistance revealed increases consistent with antagonism of channels active at baseline membrane potentials by the respective drugs (Dunn’s, control vs 4-AP p = 0.002, control vs. TEA p = 0.001, 4-AP vs TEA p >0.99).

### Total voltage-dependent K^+^ current in AVPV kisspeptin neurons has multiple components

In nocturnal rodents, positive feedback needed to generate the LH surge occurs during the afternoon of proestrus (Brown-Grant et al., 1970; Sarkar et al., 1976). During this time, both overall firing rate and burst firing of AVPV kisspeptin neurons increase relative to diestrus (Wang et al., 2016). Given the importance of K^+^ currents in determining the excitability and firing patterns of neurons (Coetzee et al., 1999; Hille, 2001), we hypothesized voltage-gated potassium currents are targets of estradiol feedback in AVPV kisspeptin neurons; we tested this using whole-cell voltage-clamp. Figure 1A shows a representative recording of total voltage-dependent K^+^ current activation. Examining this family of traces from the most hyperpolarized to the most depolarized command potentials revealed induction of one component with faster inactivation at more hyperpolarized potentials that merged with a component that activated at more depolarized potentials and only slowly inactivated. The slow kinetics of the macroscopic current resembled a K+ current previously observed in dorsal root ganglion neurons (Gold et al., 1996), and led us to extend the duration of the prepulses beyond what is used in a typical inactivation protocol to 10 s (Figure 1B). As described above, a fast component can be seen to activate and inactivate within ∼50 ms at more hyperpolarized potentials during the prepulse (expanded in Figure 1C, left); this component is prominent as a spike during the test pulse following hyperpolarized prepulses (expanded in Figure 1C, right). A second component required greater depolarization to begin activating and inactivated slowly only at more depolarized potentials. The peak total current density (derived from the voltage-clamp protocol in Figure 1A) was not different between diestrus and proestrus at any of the potentials tested (Figure 1D, two-way repeated-measures ANOVA for effect of cycle stage; Table 2). These observations, however, do not preclude latent changes attributable to individual components being differentially regulated.

**Figure 1.**
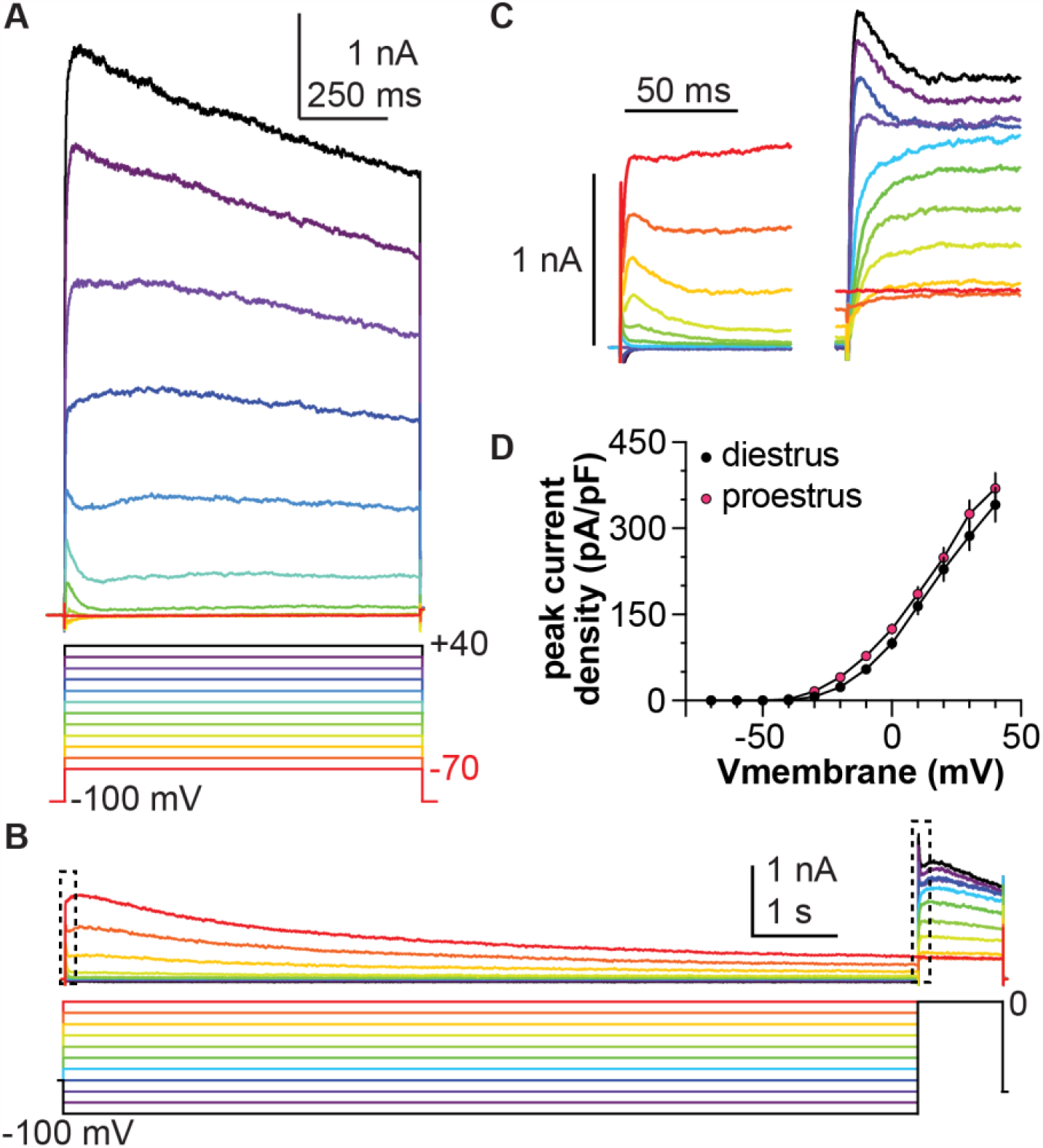
Total voltage-dependent K^+^ current in AVPV kisspeptin neurons has three distinct components. A. Representative total K^+^ current in response to the voltage-clamp protocol shown. B. Representative recording of total K^+^ current from a different cell in response to the voltage protocol shown; both a slowly-inactivating and residual (sustained) component are evident during the extended prepulse. C. Expansions of the areas within the dashed boxes in B, showing the change in inactivation rates as more depolarizing prepulses are applied (left), and how this affects both activation and inactivation during the test pulse (right). D. Mean±SEM peak current density in cells from mice in diestrus (black symbols) and proestrus (magenta symbols). Error bars are smaller than symbols for some values.

### Pharmacological treatments to isolate current components

To attempt to isolate the components of the total potassium current, recordings were made before and during application of the potassium channel blockers TEA (20 mM) or 4-AP (5 mM). TEA reduced the amplitude of both the fast and slow components, but the reduction in the slow component was proportionally greater, making the fast peak appear more prominent (Figure 2A). Lower concentrations of TEA (500 μM to 10 mM) did not block most of the slow peak amplitude and were thus less effective in revealing the fast component. The fast-transient component was markedly reduced by 5 mM 4-AP (Figure 2B, note right-shift of peak in red trace). Lower concentrations of 4-AP (100 µM to 1 mM) did not block the fast initial peak at -10 mV. These changes were attributable to drug action rather than rundown as the current remained stable in the absence of treatments over the ten minutes required for these recordings (Figure 2C).

**Figure 2.**
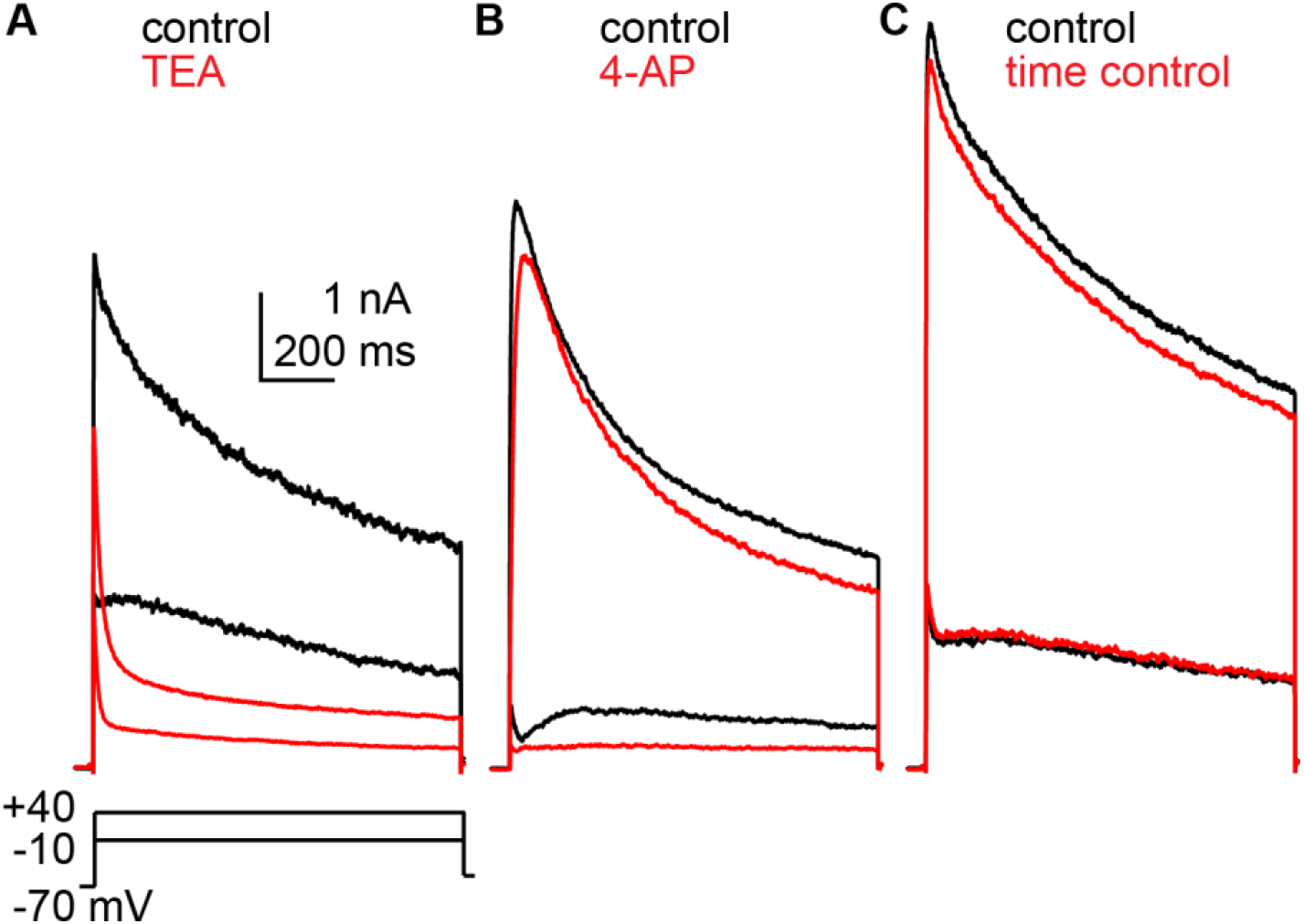
Potassium current recorded from three different cells before and during application of potassium channel blockers. A. 20 mM TEA. B. 5 mM 4-AP. C. time control.

Individual components of the total current may have distinct roles in determining firing output and may also be independently regulated by cycle stage. The differential effects of TEA and 4-AP were utilized to perform more detailed characterization of the sub-components of the total potassium current in AVPV neurons in slices from diestrous and proestrous mice. To increase recording yield, we switched from a within-cell design to a population (i.e., among-cell) design, with drug present from the time the slice was placed in the recording chamber. In these population studies, the longer exposure to TEA caused a larger reduction in peak current density. Specifically, the peak current density at +40 mV during TEA treatment was 76.6 ± 5.1% of pretreatment values in within-cell experiments, but only 53.0 ± 3.7% of untreated cells in population studies. This greater reduction was not observed with 4-AP (the peak current density during 4-AP treatment was 76.6 ± 4.9% of the control period for within-cell studies (n=3), and 76.1% of untreated controls in population studies (n=4)).

### The slow transient component is increased on diestrus

The extended duration voltage-clamp protocols were used to characterize the slow-transient potassium current in the presence of 4-AP; representative traces are shown in Figure 3. Peak current density was greater in cells from diestrous (n=12) than proestrous (n=13) mice at more depolarized potentials (Figure 3C, two-way repeated-measures ANOVA for effect of cycle stage; Table 2). The slow current was activated at command voltages at and depolarized relative to -20 mV. Normalized activation and inactivation curves were not different between groups (Figure 3D, two-tailed Student’s *t*-test with Welch’s correction for V50 activation, inactivation; Table 3). Timecourse of inactivation was not different between diestrus and proestrus (Figure 3F, two-way repeated-measures ANOVA for effect of cycle stage; Table 2). In contrast, recovery from inactivation was more rapid during proestrus (Figures 3G, H, two-way repeated-measures ANOVA for effect of cycle stage; Table 2). In recovery from inactivation experiments, the recovery of two currents was evident. The fast transient was visible when the hyperpolarizing recovery prepulse was shorter than 200 ms (Figure 3G, arrow). This may be attributable to the voltage-dependence of 4-AP action, in which the channel becomes unblocked during depolarization (Choquet and Korn, 1992; Kehl, 2017). The slow current was visible 70-150 ms into the test pulse. To avoid contamination with the fast current, for this protocol only the peak of the slow transient current was taken as the maximum current 70-150 ms after the start of the test pulse.

**Table 2.** Two-way ANOVA analyses of K^+^ current properties. Bold font indicates p<0.05.

|  | n cells, n animals |  | estrous stage | estrous stage x Vm or duration interaction |
| --- | --- | --- | --- | --- |
| TOTAL | di | pro |  |  |
| density | 10, 4 | 17, 5 | Diff, -15.42 [CI -40.49, 9.660]<br>F(1, 25) = 1.603; p=0.217 | F (11, 275) = 0.6496<br>p=0.785 |
| **SLOW** |  |  |  |  |
| density | 12, 5 | 13, 5 | Diff, 24.07 [CI -5.523, 53.66]<br>F (1, 23) = 6.503; p=0.106 | **F (8, 184) = 10.03**<br>**p<0.001** |
| activation | 12, 5 | 13, 5 | Diff, -0.01901 [CI -0.06921, 0.03120]<br>F (1, 25) = 5.283; p=0.442 | F (11, 275) = 3.336<br>p=0.169 |
| inactivation | 9, 4 | 12, 5 | Diff, -0.0009547 [CI -0.06269, 0.06078]<br>F(1, 20) = 0.1782; p=0.975 | F (12, 240) = 0.1988<br>p>0.999 |
| rate of inactivation | 10, 7 | 9, 4 | Diff, -0.03472 [CI -0.08026, 0.01082]<br>F (1, 17) = 2.587; p=0.126 | F (14, 238) = 1.336<br>p=0.187 |
| rate of recovery | 9, 5 | 13, 5 | **Diff, -0.05901 [CI -0.09461, -0.02342]**<br>**F (1, 20) = 10.52; p=0.002** | **F (14, 280) = 4.006**<br>**p<0.001** |
| **FAST** |  |  |  |  |
| density | 11, 4 | 13, 4 | Diff, -6.607 [CI -22.53, 9.317]<br>F (1, 22) = 0.7404; p=0.399 | F (11, 242) = 0.9366<br>p=0.506 |
| activation | 11, 4 | 13, 4 | Diff, 0.01876 [CI -0.03188, 0.06940]<br>F (1, 23) = 0.6133; p=0.450 | F (8, 184) = 1.475<br>p=0.981 |
| inactivation | 11, 4 | 12, 4 | **Diff, -0.05682 [CI -0.1119, -0.001691]**<br>**F (1, 21) = 4.594; p=0.044** | **F (9, 189) = 4.041**<br>**p<0.001** |
| rate of inactivation | 10, 4 | 12, 4 | Diff, -0.0085 [CI -0.03704, 0.02004]<br>F (1, 20) = 0.3859; p=0.541 | F (11, 220) = 0.4253<br>p=0.944 |
| rate of recovery | 10, 4 | 12, 4 | Diff, -0.01081 [CI -0.03695, 0.01532]<br>F (1, 20) = 0.7448; p=0.398 | F (11, 220) = 0.3670<br>p=0.967 |
| **RESIDUAL** |  |  |  |  |
| density | 11, 4 | 13, 4 | Diff, 0.07334 [CI -3.263, 3.409]<br>F (1, 22) = 0.002079; p=0.964 | F (11, 242) = 0.05676<br>p>0.999 |
| activation | 11, 4 | 13, 4 | Diff, 0.01068 [CI -0.03932, 0.06068]<br>F (1, 22) = 0.1962; p=0.662 | F (11, 242) = 0.2095<br>p=0.997 |

**Table 3.**
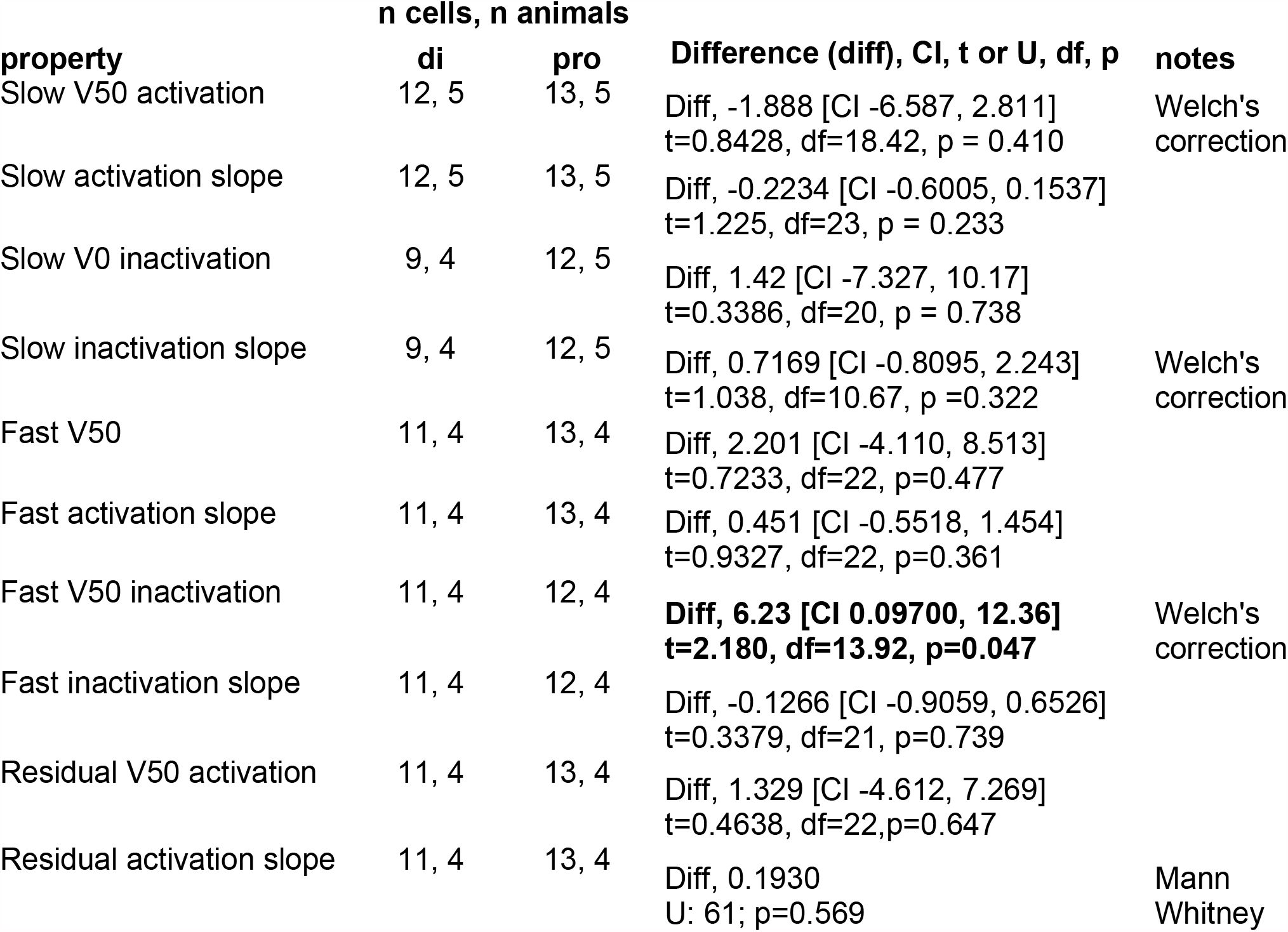
Two-sample analyses of K^+^ current properties. Bold font indicates p<0.05. Differences shown for means for normally-distributed data and medians for non-normally-distributed data.

**Figure 3.**
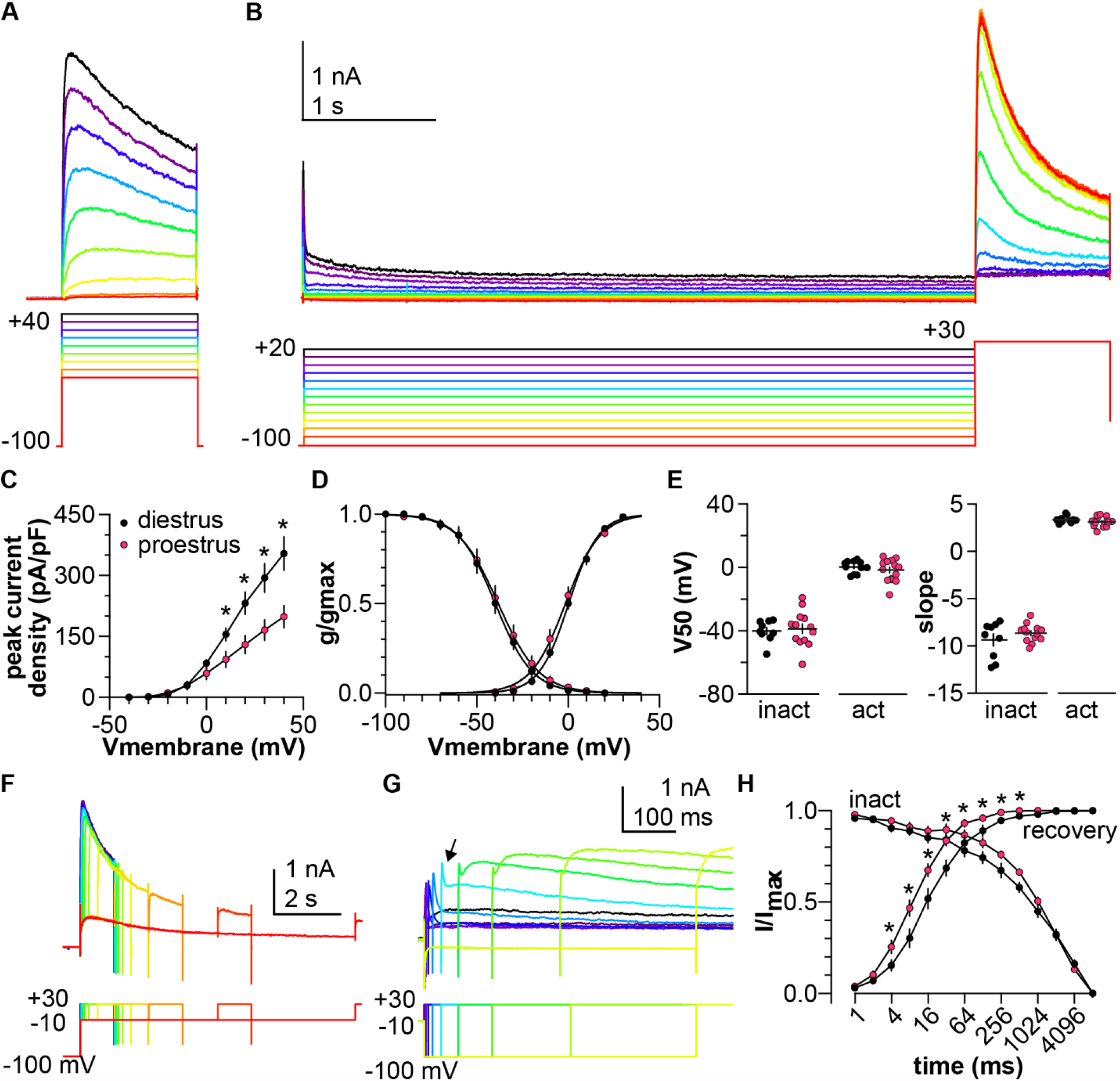
4-AP-resistant slow-transient voltage-dependent K^+^ current is larger on diestrus. A. Representative traces (top) in response to the activation voltage-clamp protocol (bottom) in the presence of 5 mM 4-AP. B. Representative traces (top) in response to the inactivation voltage-clamp protocol (bottom). Offline leak subtraction was applied to test pulses only, hence the capacitance transient is visible at the start of the recording. C. Mean ± SEM peak current density from cells in mice in diestrus (black symbols) and proestrus (magenta symbols). D. Mean ± SEM normalized conductance. Solid lines are Boltzmann fits to the mean data. E. Individual values and mean ± SEM parameters obtained from Boltzmann fits to normalized conductance curves for each cell. F, G. Representative traces (top) in response to the voltage-clamp protocol (bottom) used to measure the time-dependence of inactivation (F) and recovery (G). Arrow denotes peak of fast transient current. H. Mean ± SEM normalized peak current vs. prepulse duration for time-dependence of inactivation (inact) and recovery from inactivation. Error bars are smaller than symbols for some values. * p<0.05

### The inactivation curve of the fast-transient component is depolarized during proestrus

Bath-applied 20 mM TEA preferentially reduced the amplitude of the slow-transient current, allowing better characterization of the fast-transient current. Because this fast-transient component activated and inactivated quickly, voltage-clamp protocols with more traditional durations were utilized (Figure 4A). Under these recording conditions, a fast-transient current began to activate between -50 mV and -40 mV (Figure 4C); current reached a non-zero steady state by the end of the test pulse, suggesting a contribution by non- or slowly-inactivating channels. To isolate the fast-transient current from this non-inactivating residual component, a voltage-based subtraction method was used. Subtraction of the residual current following a depolarizing prepulse (Figure 4A, middle) yielded a transient current that was almost fully inactivated by the end of the test-pulse (Figure 4A, right). The V50 activation of the residual current was more depolarized than that of the fast-transient current at both cycle stages (Figure 4B-D diestrus n=11, proestrus n=13, one-way ANOVA F(3,44) =10.8, p<0.001; Tukey residual vs fast p=0.001 for proestrus, residual vs fast p=0.002 for diestrus). There was no effect of cycle stage on the peak current density of either the fast-transient or residual component (Figure 4B, two-way repeated-measures ANOVA for effect of cycle stage; Table 2). In contrast, cycle stage affected voltage dependence of the fast-transient component. Specifically, inactivation of the fast transient occurred at more depolarized potentials on proestrus than diestrus (Figure 4C,E, two-way repeated-measures ANOVA for effect of cycle stage on inactivation; Table 2; unpaired two-tailed Student’s *t*-test with Welch’s correction of V50 inactivation; Table 3), Cycle stage did not, however, affect the voltage-dependence of activation (Figure 4C, two-way repeated-measures ANOVA for effect of cycle stage on activation curve; Table 2) or V50 activation (Figure 4D, unpaired two-tailed Student’s *t*-test; Table 3). Full inactivation and full recovery of the fast transient occurred in less than 100 ms (Figure 4F, G). There was no group difference in either of these properties (two-way repeated-measures ANOVAs for effect of cycle stage; Table 2).

**Figure 4.**
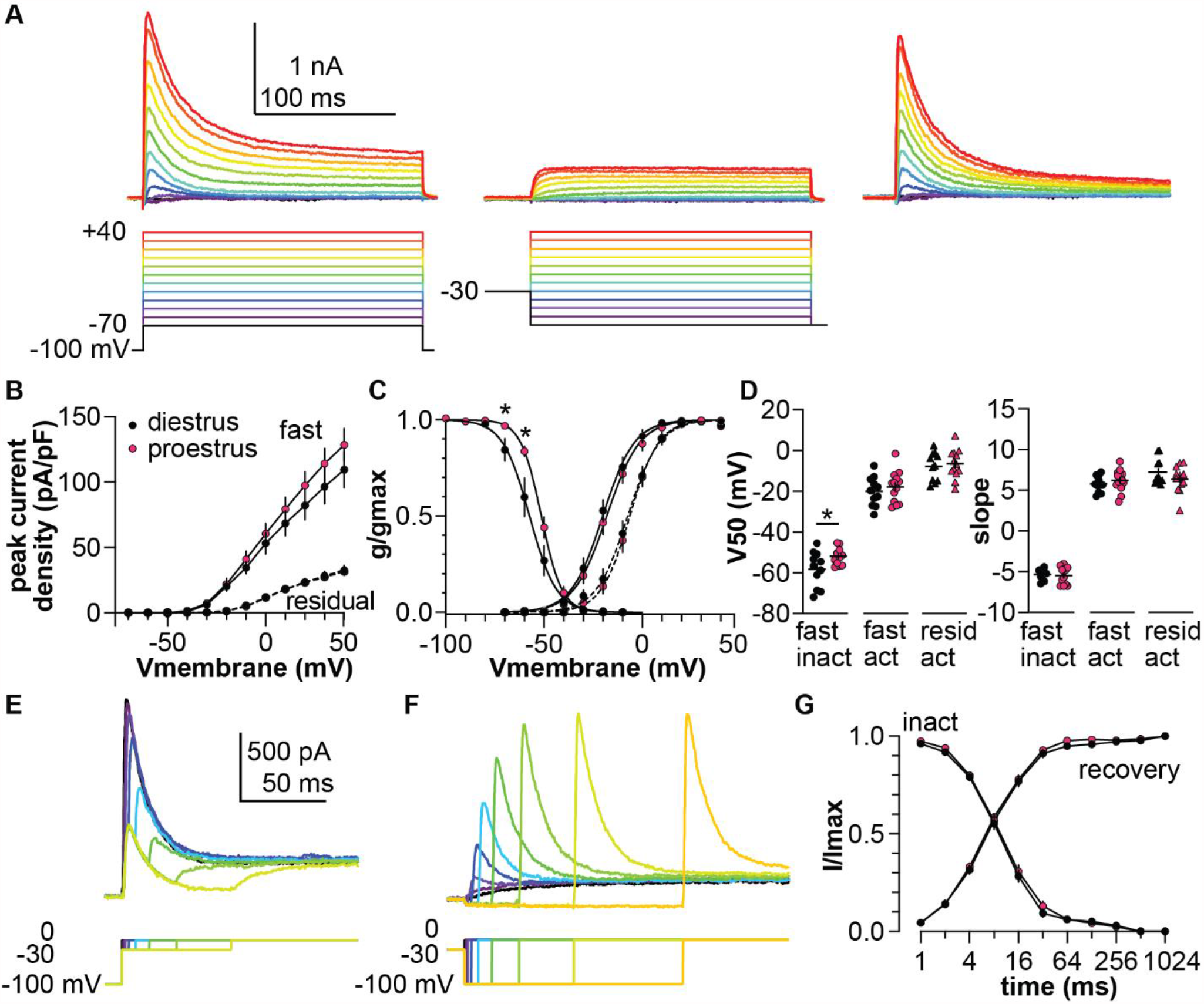
TEA-resistant fast-transient voltage-dependent K^+^ current has a depolarized inactivation curve during proestrus. A. Left: Representative unsubtracted K^+^ current (top) in response to the voltage-clamp protocol (bottom) in presence of 20 mM TEA. Middle: Residual current in the same cell with a -30 mV prepulse. Right: Fast transient current yielded by subtracting residual current from raw current. B. Mean ± SEM peak current density in cells from mice in diestrus (black symbols) and proestrus (magenta symbols). C. Mean ± SEM normalized conductance. Solid lines represent Boltzmann sigmoidal fits to the mean data for the fast transient, dash lines the fits for the residual current. D. Individual values and mean ± SEM parameters obtained from Boltzmann fits to normalized conductance curves for each cell. E,F. Representative traces (top) in response to the voltage-clamp protocol (bottom) used to measure the time-dependence of inactivation (E) and recovery (F). G. Mean ± SEM normalized peak current vs. prepulse duration for time-dependence of inactivation (inact) and recovery from inactivation. Error bars are smaller than symbols for some values. * p<0.05

### Modeling conductances

Computational modeling was used to better understand the role of estrous cycle modulation of multiple conductance types in the control of AVPV kisspeptin neuron firing. Specifically, we examined if estrous cycle-related shifts in K^+^ current components and/or subthreshold depolarizing currents alter the firing dynamics of these cells. To model subcomponents *in silico*, Hodgkin-Huxley conductance models were fit to mean ± SEM recordings of drug- and voltage-separated subcomponents of the K^+^ current: slow, fast and residual. This involved adding individual subcomponents to single-compartment neuron models and running such models through the same voltage-clamp protocols used experimentally, generating membrane current over time plots. Model output was compared to experimental data (Figure 5), and m_∞_(V), h_∞_(V), τ_m_(V), and τ_h_(V) function parameters were changed iteratively to improve the quality of fit. Steady-state activation and inactivation curves and I(density)-V plots were calculated from the model output and compared to experimental data for K^+^ currents (Figure 6A, B, D) and for CaT (Figure 6C, F) and NaP (Figure 6E) currents reported (Zhang et al., 2015; Wang et al., 2016).

**Figure 5.**
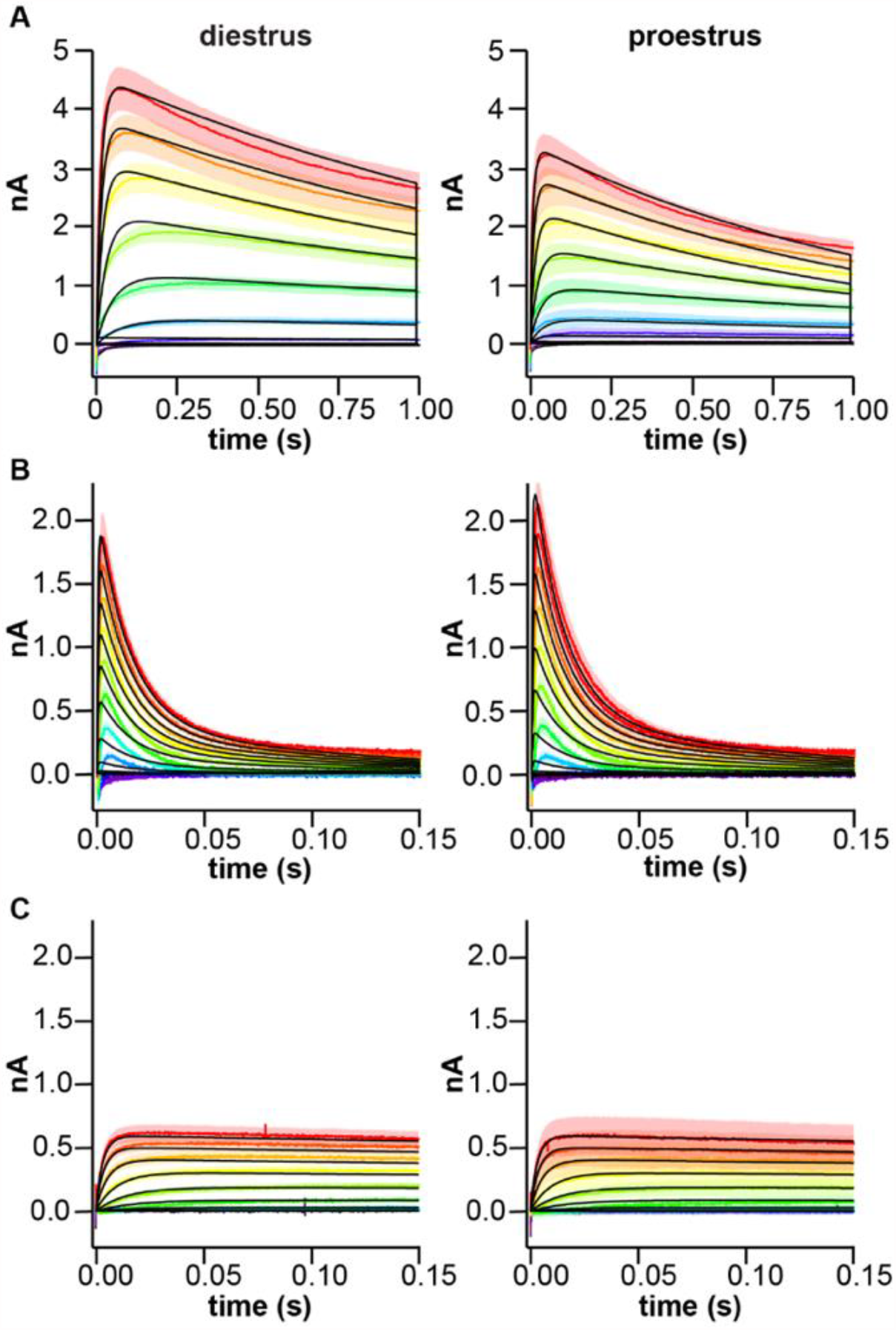
Potassium conductance model output vs experimental data for voltage steps. Rainbow colors indicate mean ± SEM current traces recorded at different test potentials; black lines show model simulations. In each panel, diestrus is on the left and proestrus on the right. A. Slow (voltage protocol as in Figure 3A). B. Fast (voltage protocol as in Figure 4A). C. Residual (voltage protocol as in Figure 4A).

**Figure 6.**
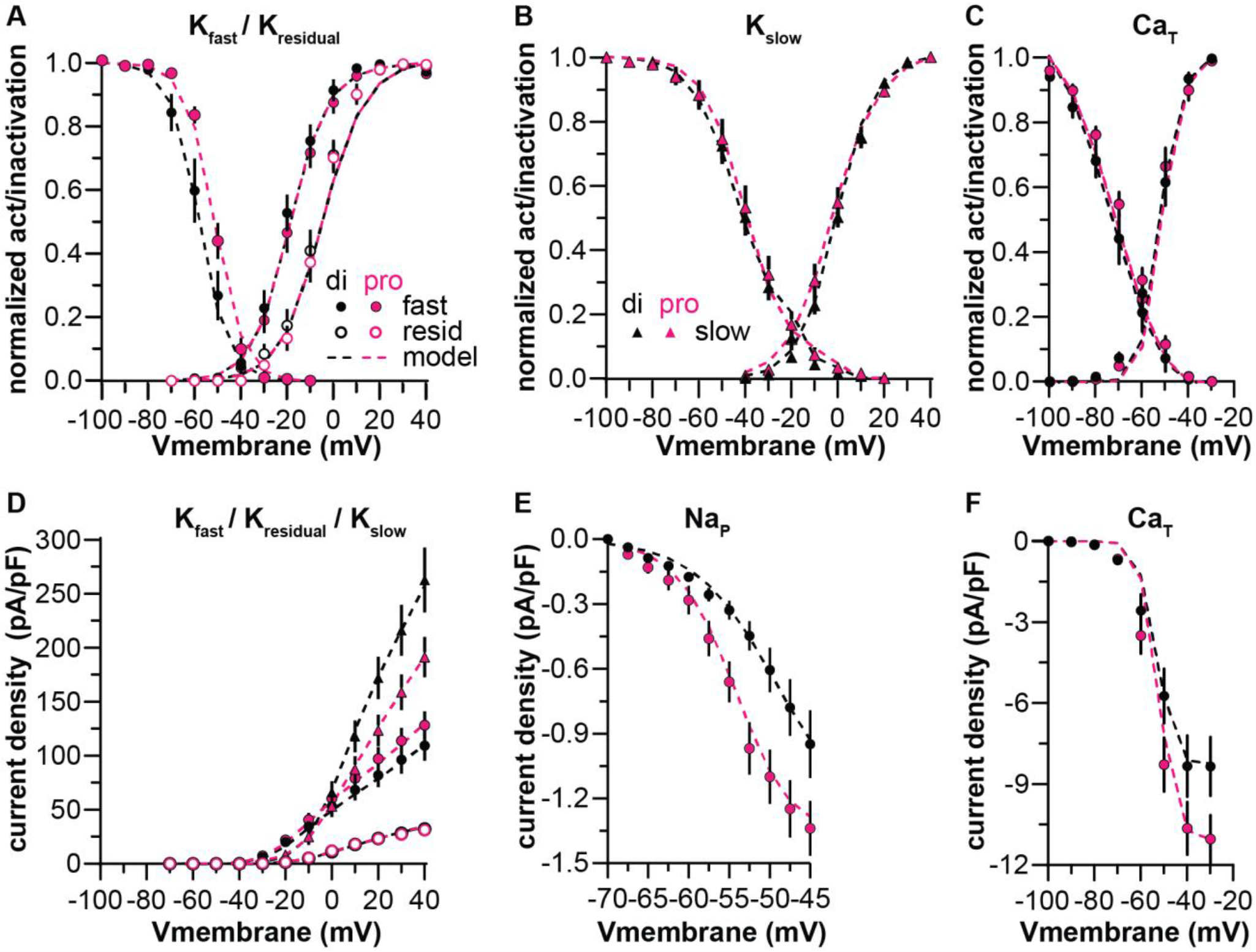
Conductance models vs data for steady state activation/inactivation and current density experimental data. A-C. Steady state activation/inactivation curves calculated from voltage-clamp simulations (dashed lines) compared to corresponding mean ± SEM experimental data (symbols) from diestrous (black) and proestrus (magenta) groups. D-F. Peak current density for various conductances. Experimental data for K^+^ current subcomponents are the same as in Figures 3 and 4 and replotted here for ease of comparison. Data points for NaP (E) and CaT (C, F) are adapted from Wang et al., 2016.

To reconstruct the total K^+^ current from the sum of the subcomponents, the set of I_slow_, I_fast_, and I_resid_ currents for each group were included in single neuron models, and the models were run through the same voltage-clamp protocol used for recordings in Figure 1A. Initial simulations indicated the sum of fast, slow, and residual currents as fit to their isolated recordings was not enough to account for the full amplitude of the total K^+^ current, suggesting subcomponents were blocked to some degree by the drug used to isolate them (i.e., 4-AP blocked a portion of the slow current, TEA blocked some of the fast and residual currents). To overcome this problem, particle-swarm optimization was used to minimize the error between the model-generated current trace and the mean ± SEM trace by optimizing 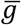 parameters of each subcomponent, with equal weighting across the trace. This preserved the voltage-sensitivity and kinetics of the subcomponents as fit in the drug- and voltage-separated recordings (Figure 5) while providing a better fit to the experimental data. Parameter values are shown in Tables 4-5. This approach provided a good fit to peak current density (Figure 7A) and the first ∼250 ms of the total K^+^ current (Figure 7B). The error between the model and data increased as the simulation progressed, however. This error could be reduced by decreasing the inactivation rate of I_slow_, but this modification had no effect on firing behavior as measured in response to 500 ms current pulses, suggesting this shift in kinetics does not substantially influence firing output at these timescales (not shown).

**Table 4.**
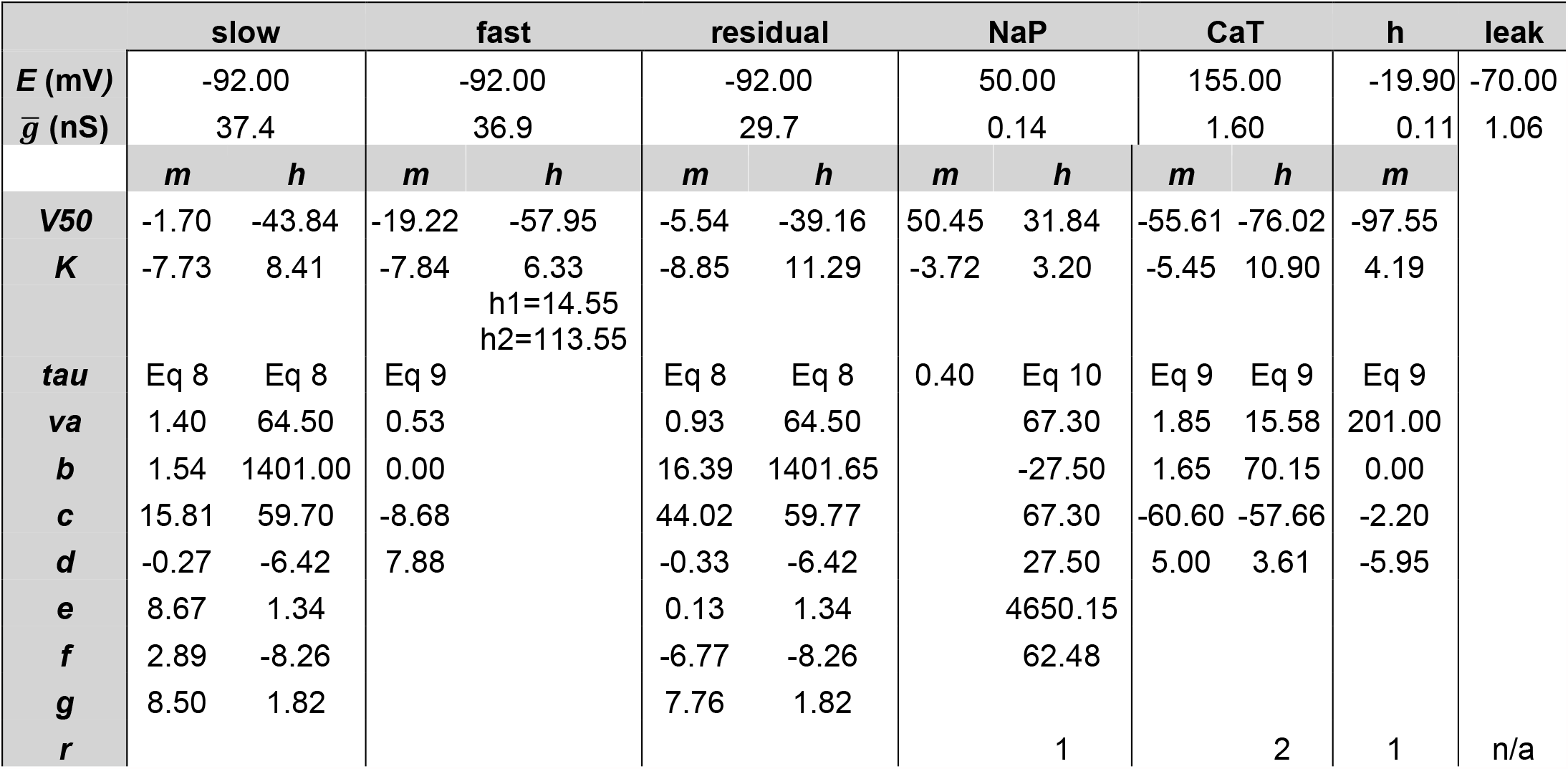
Model parameters for diestrus

**Table 5.**
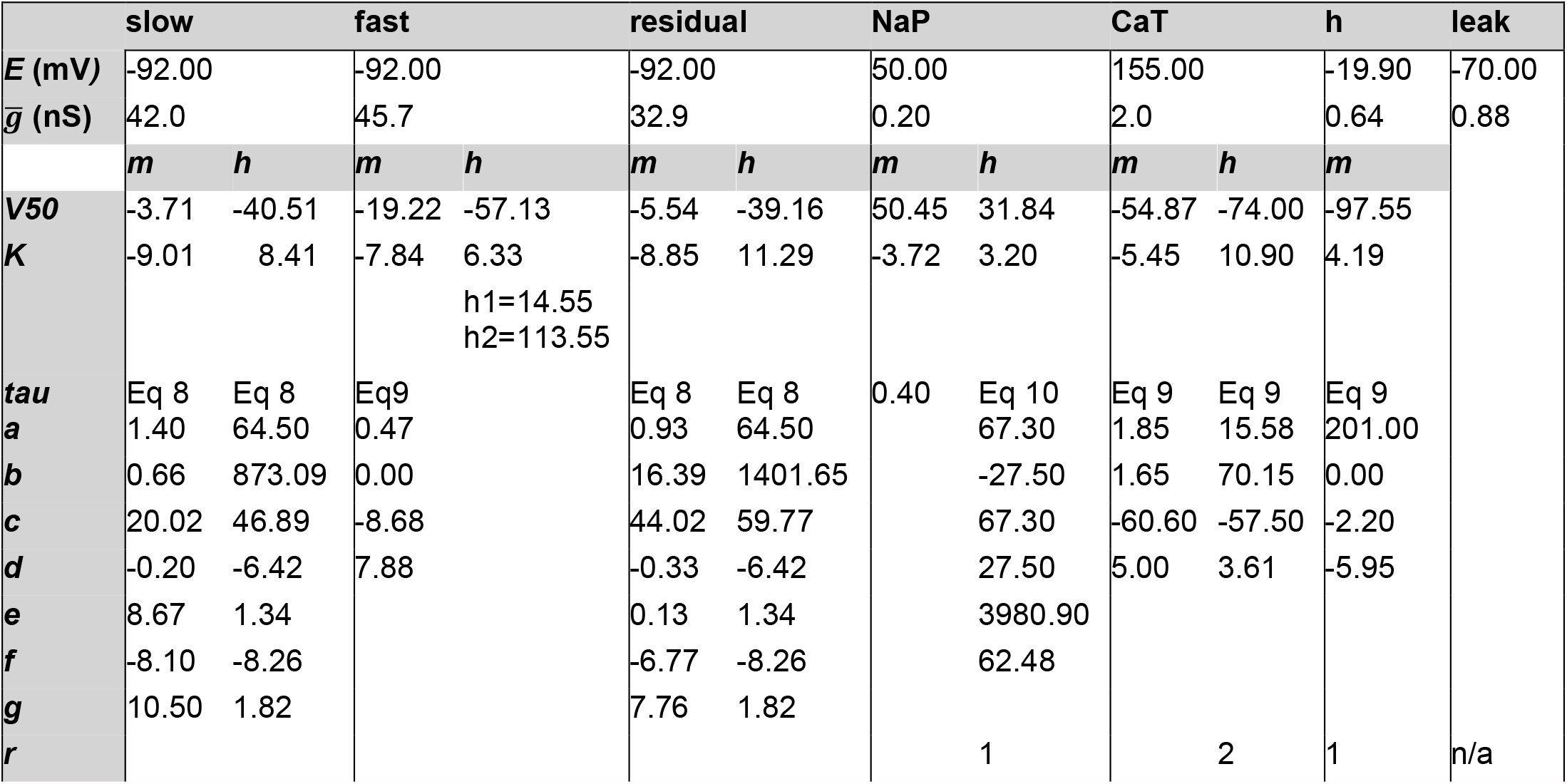
Model parameters for proestrus

**Figure 7.**
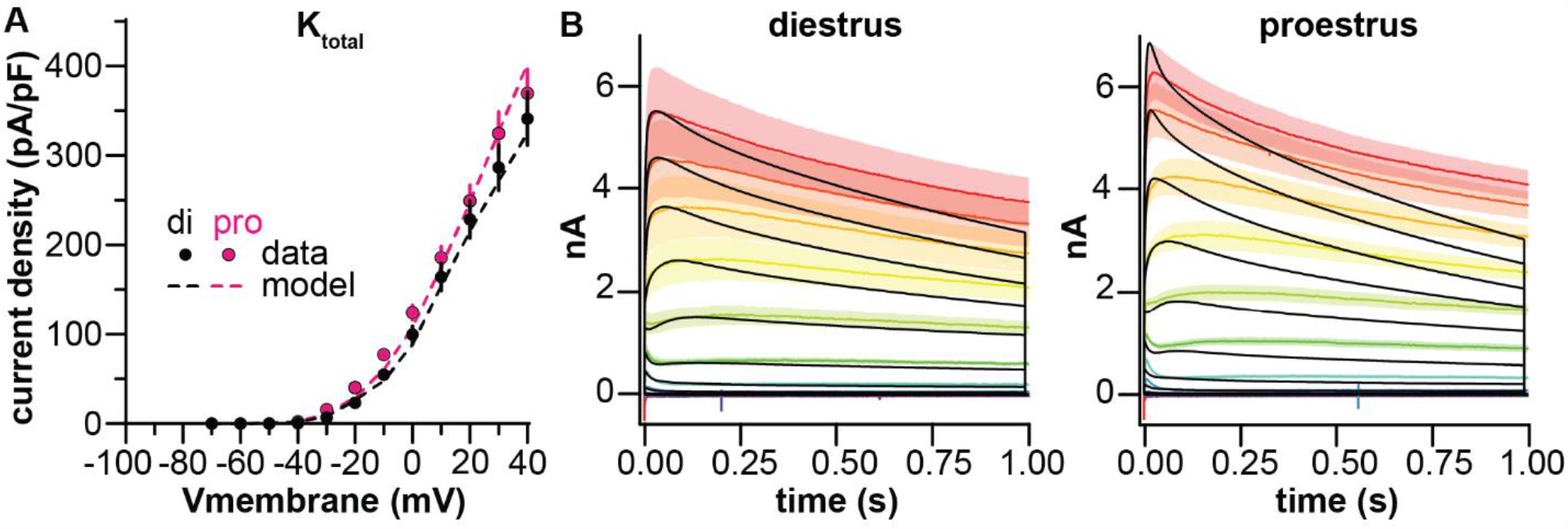
Reconstruction of the total K^+^ current from the sum of the three subcomponents. A. Peak current density when I_fast_, I_slow_, and I_resid_ are simulated together in the same model (dashed lines) after 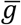 optimization to correct for suppression by TEA/4-AP. Mean ± SEM symbols (black: diestrus, magenta: proestrus) and voltage-clamp protocols are the same as shown in Figure 1A and D and are reproduced for ease of comparison. B. Mean ± SEM experimental current traces at different test pulses (rainbow colors) and model simulation (black).

### Simulations of firing

K^+^ conductance models from fits to the total K^+^ current, CaT, NaP, and HCN conductances for the diestrous and proestrous groups were combined into single-compartment neuron models, thus creating neuron models with seven voltage-gated conductances and one leak conductance each. A transient sodium (NaT) conductance is needed to enable action-potential firing and was thus added to the models at this stage (Table 6). Transient sodium currents of AVPV kisspeptin neurons have not been measured. We thus estimated NaT parameters using action potential properties such as threshold and peak amplitude. NaT was kept identical between diestrus and proestrus as these action potential properties did not change with cycle stage previously (Wang et al., 2016) or this study (threshold: diestrus -51.8 ± 1.0 mV, n = 12 cells from 5 animals, proestrus -51.6 ± 0.6 mV, n = 17 cells from 6 animals, two-tailed, unpaired Student’s *t*-test, p = 0.85, t=0.1906, df=27, amplitude diestrus 82.1 ± 1.9 mV, n = 12, proestrus 80.5 ± 1.1 mV, n = 17, two-tailed unpaired Student’s *t*-test, p = 0.4501, t=0.72, df=22) suggesting estrous cycle modulation of this conductance is minimal. Using the same NaT conductance in the diestrous and proestrous models generated action potential shapes similar to experimental action potentials (Figure 8C). The diestrous and proestrous models were tested with the same current injection protocols used in recordings and demonstrated F-I curves similar to the mean ± SEM data, indicating that the models could reproduce the firing behavior of AVPV kisspeptin neurons recorded in brain slices (Figure 8D).

**Table 6.**
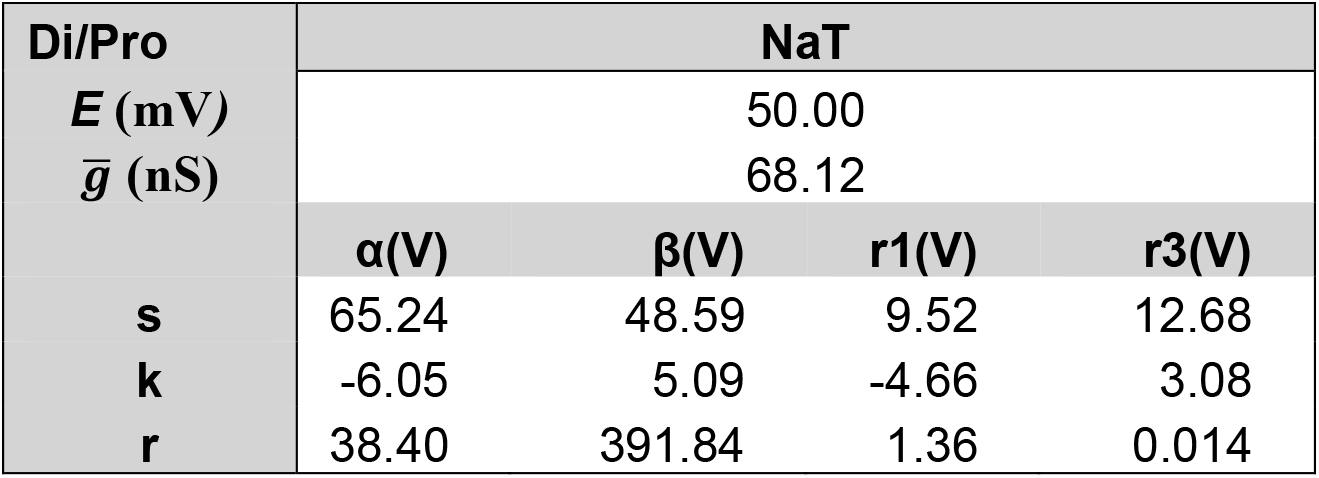
NaT parameters

**Figure 8.**
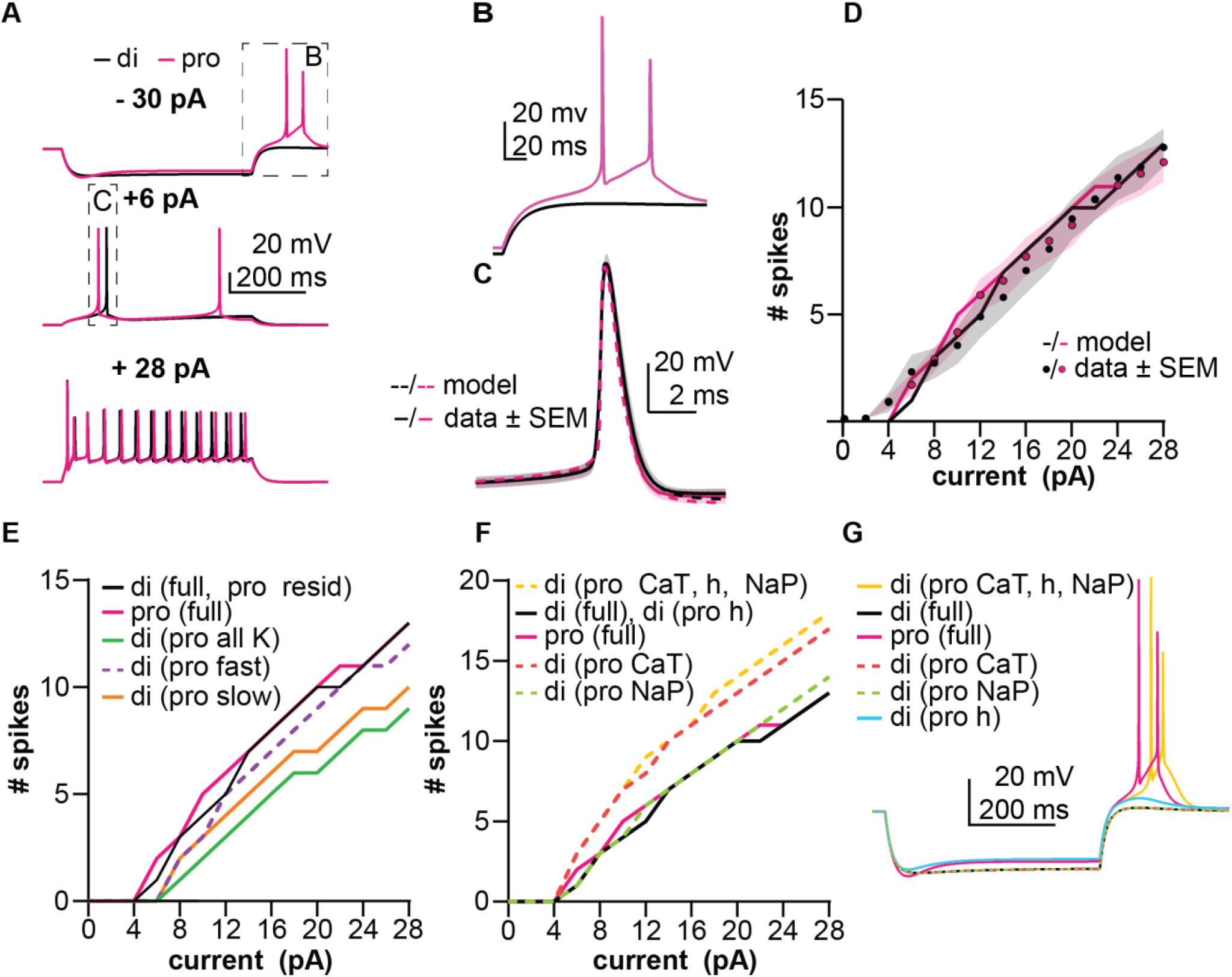
Simulations of firing from a baseline of -70 mV. A. Performance of diestrous (black) and proestrous (magenta) models in response to -30 pA (top), +6 pA (middle), and +28 pA (bottom) applied current. Boxed regions are shown with expanded axes in the indicated subfigure. B. Post-hyperpolarization rebound of diestrous and proestrus models from subfigure A. C). Rheobase action potentials for both models (dashed lines) from A (middle) and experimental counterparts (solid lines, mean ± SEM). D. Firing vs current (F-I) curves for diestrous and proestrous models (lines) and experimental counterparts (circles are means, shading is SEM). E. F-I performance of hybrid models in which one or multiple K^+^ conductances in the diestrous model was substituted for a proestrous counterpart. Non-hybrid models from part D (di full and pro full) are reproduced here to facilitate comparison in this panel as well as F and G. F. F-I performance of hybrid models in which one or multiple subthreshold depolarizing currents in the diestrous model were replaced with proestrous counterparts. G. Rebound bursting performance of hybrid models in response to -30 pA current injection.

These models also faithfully reproduced cycle-dependent differences in post-hyperpolarization rebound firing behavior observed (Wang et al., 2016). Specifically, cessation of hyperpolarization initiated rebound spikes in the proestrous but not the diestrous model (Figure 8B). To determine the basis for this difference as well as to further understand how each conductance influences firing behavior, we performed a series of simulations involving swapping individual or multiple conductances from one model into the other. In the diestrous model, replacing all K^+^ conductances with their proestrus counterparts had a suppressive effect, depolarizing the rheobase and reducing the number of spikes fired in response to all depolarizing stimuli (Figure 8E). Of the three potassium current subcomponents, proestrous I_slow_ was the most suppressive when individually substituted into the diestrous model (Figure 8E). The diestrous model with only I_fast_ substituted for the proestrous counterpart had a depolarized rheobase but the F-I curve (Figure 8E) had a similar slope. Changing from diestrous to proestrus I_residual_ did not change firing behavior (Figure 8E), consistent with this current being similar between groups (Figure 4). Of the subthreshold depolarizing conductances, proestrous CaT caused the greatest increase in excitability when substituted into the diestrous model, generating more spikes at every depolarizing stimulus after rheobase was achieved, but not changing rheobase itself (Figure 8F). Rebound firing only occurred in the diestrous model when all three subthreshold depolarizing currents were replaced with their proestrous counterparts (Figure 8F, G). This model, which had diestrous K^+^ conductances and proestrus CaT, NaP, and h-current, had the greatest excitability of any model tested. It is important to point out that changes in excitability were not attributable to differences in either baseline membrane potential or input resistance as these were the same among models. Comparing this hybrid model to the proestrus-only model thus further indicates the suppressive effect of proestrous K^+^ currents.

## Discussion

AVPV kisspeptin-expressing neurons are key mediators of estradiol positive feedback–a process critical for ovulation. We studied intrinsic properties of these neurons to determine how they become more active during proestrus, the phase of the rodent estrous cycle when positive feedback culminates in the ovulation-inducing GnRH/LH surge. AVPV kisspeptin neurons exhibit three types of voltage-gated potassium currents; two were modulated in an estrous-cycle dependent manner. Unexpectedly, the effects of the proestrous K^+^ currents were suppressive but countered by cycle-dependent shifts in inward currents active in the subthreshold range. The proestrous shift in K^+^ currents did not, however, block increased rebound firing enabled by the proestrous shift in inward currents.

This finding that the K^+^ currents on proestrus were suppressive was initially surprising given substantial evidence indicating AVPV kisspeptin neurons are activated during estradiol positive feedback (Zhang et al., 2015; Wang et al., 2016, 2019). Much of this evidence was gathered using experimental models of estradiol feedback involving ovariectomy plus estradiol replacement (OVX+E) as a constant-release implant producing daily surges (Christian et al., 2005), or an implant followed several days later by an estradiol benzoate injection that triggers an LH surge the next day (Bronson and Vom Saal, 1979). These models allow study of estradiol feedback in isolation of other ovarian factors. In both models, AVPV kisspeptin neurons have increased c-fos expression during the LH surge (Adachi et al., 2007; Porteous and Herbison, 2019). Increases in spontaneous firing rate during positive feedback in the daily surge model are similar to those observed in naturally cycling animals on proestrus (Wang et al., 2016, 2019). Interestingly, given the changes in spontaneous firing, no studies have reported that feedback or estrous cycle stage affect the F-I curve of AVPV kisspeptin neurons. Our data and simulations suggest this lack of effect on the F-I curve is attributable to more suppressive K^+^ currents during proestrus counteracting larger inward currents during depolarization, without inhibiting rebound burst firing.

Estradiol feedback also affects voltage-gated K^+^ currents in other central neurons controlling reproduction. In GnRH neurons, inactivation of the transient K^+^ current is depolarized by positive feedback in the daily surge model to a similar degree observed in the present study (DeFazio et al., 2002). In contrast to AVPV neurons, however, peak K^+^ current densities in GnRH neurons are reduced during positive feedback, and excitability is increased (Adams et al., 2018).

Interestingly, positive feedback effects in GnRH neurons may be at least in part attributable to increased kisspeptin receptor activation, as kisspeptin treatment has similar effects on transient K^+^ currents in these cells (Pielecka-Fortuna et al., 2011). In arcuate kisspeptin neurons, K^+^ currents were compared between open-loop (OVX) and estradiol negative feedback (DeFazio et al., 2019), with negative feedback reducing peak transient and sustained K^+^ currents compared to open-loop but not affecting voltage sensitivity of activation/inactivation. Together these observations suggest the effects of estradiol and cycle stage on voltage-gated K^+^ currents in the reproductive neuroendocrine circuitry depend on both cell type and type of feedback involved.

The subcellular mechanisms linking estradiol feedback to these effects have not been extensively studied but may include changes in channel subunit expression/composition, subcellular location of channels, binding by partner proteins, and/or subunit phosphorylation state (Coetzee et al., 1999; Levitan, 2006). In GnRH neurons, estradiol or cycle stage regulates expression of Ca^2+^, HCN, and SK channel genes (Zhang et al., 2009; Bosch et al., 2013; Rønnekleiv et al., 2015; Vastagh et al., 2019). Not all these effects are specific to GnRH neurons, as K^+^ channel genes are estradiol-sensitive in the paraventricular and arcuate nuclei (Qiu et al., 2006; Roepke et al., 2007; Lee et al., 2013; Yang et al., 2016). Estradiol affects transcription of ion channels outside the central nervous system in neural (Du et al., 2014) and non-neural tissue (Marni et al., 2009; Banciu et al., 2018). In addition to transcription, estradiol action through membrane-associated signaling pathways can rapidly modulate L- and R-type Ca^2+^ and K-ATP currents (Sun et al., 2010; Zhang et al., 2010). Estradiol positive feedback effects on K^+^ currents in GnRH neurons are attenuated by a broad-spectrum kinase inhibitor, suggesting a role of phosphorylation (DeFazio and Moenter, 2002). Estradiol may also interact directly with channels, as with voltage-gated BK channels in vascular smooth muscle (Granados et al., 2019). The effects of estradiol on CaT currents in AVPV kisspeptin neurons, however, are dependent on actions via ERα as cre-lox or CRISPR mediated ERα knockdown in these cells eliminates estradiol effects (Wang et al., 2019).

The present results add important information to a growing list of voltage-gated conductances in AVPV kisspeptin neurons that are modified by the estrous cycle (Piet et al., 2013; Zhang et al., 2015; Wang et al., 2016). We used *in silico* approaches to amalgamate multiple experimental findings into a more comprehensive understanding of how hormonal manipulations/estrous cycle shape the membrane response of these cells as in other reproductive neuroendocrine neurons (Moran et al., 2016; Adams et al., 2018; Mendonça et al., 2018; DeFazio et al., 2019). Simulation results suggest modulation of K^+^ currents across the negative to positive feedback transition do not increase AVPV neuron excitability or enable rebound burst firing, as had been suggested by blocking the fast-transient current with 4AP in current-clamp studies (Wang et al., 2016). Rather, proestrous shifts in subthreshold inward currents CaT, NaP, and HCN counteract suppressive K^+^ currents during proestrus and enable rebound firing. Involvement of the former currents in rebound firing is consistent with their role in shaping firing behavior (Lüthi and McCormick, 1998; Bevan and Wilson, 1999; Hille, 2001; Perez-Reyes, 2003).

The finding that cycle-dependent changes in intrinsic properties lead to no net change in excitability begs the question of how increased firing rates during proestrus are maintained when receptors for fast glutamate and GABA transmission are blocked (Wang et al., 2016). First, neuromodulation by catecholamines, acetylcholine, and neuropeptides may occur. Second, proestrous increases in spontaneous firing rate may be due to the emergence of rebound bursting (Wang et al., 2016). The suppressive K^+^ current changes observed on proestrus did not inhibit this process and may bestow properties to the cell that are otherwise beneficial. For example, these more suppressive K^+^ currents may serve a protective/ homeostatic function by limiting the increase in excitability enabled by increases in depolarizing currents. An optogenetic study examining the relationship between AVPV neuron stimulation and GnRH neuron firing found delayed activation of GnRH neurons characteristic of kisspeptin receptor activation plateaus at 10 Hz (Piet et al., 2018), suggesting firing frequencies above this are unnecessary. Third, excitation-release coupling in AVPV neurons may be regulated by the estrous cycle. Small increases in release probability or the readily releasable pool of kisspeptin could have substantial downstream effects on GnRH neuron activation with only modest differences in AVPV neuron firing frequency. Of note, kisspeptin expression by these cells is increased during positive feedback (Smith et al., 2005; Gottsch et al., 2006). An important consideration for brain slice preparations is the disruption of neuronal projections that likely play a role in cycle-dependent changes. In rodents, input from the circadian clock is critical for surge induction (van der Beek, 1996); these and other afferents may be regulators of AVPV neuron activity *in vivo* and would not be present in the slices used (Watson et al., 1995).

*In silico* approaches provide the advantage of being able to test multiple hypotheses quickly but also have several caveats. First, while we sought to make the models reasonably comprehensive by including characterizations of currents reported in the literature, there are likely uncharacterized currents expressed in AVPV neurons that may change with cycle stage and/or influence firing behavior that were not included. Second, and related to the first point, no studies have characterized the transient sodium current in AVPV neurons. We adapted a Markov model of NaT used in a GnRH neuron model to enable firing in these models (Adams et al., 2018). Estimating this current was guided by fitting the rising phase of the action potential to experimental data, the phase of the action potential most highly influenced by sodium channel properties (Kress et al., 2010). Third, no computational model is ever a perfect representation, and it is possible that other combinations of properties for the components of this model could produce similar results in firing output. Despite these limitations, our voltage-clamp models are well fit, and the firing simulations reasonably approximated the firing behavior of these cells.

Together these results point to changes in multiple voltage-dependent currents in these cells through the reproductive cycle. Our data and firing simulations suggest estrous cycle effects on these currents are unlikely to explain the increase in AVPV kisspeptin neuron spontaneous activity during the afternoon of proestrus, as changes in typically hyperpolarizing and depolarizing currents exert reciprocal effects on excitability, though the latter contribute to the emergence of post-inhibitory rebound burst firing. These findings motivate further study of the inputs controlling firing behavior of these cells as well as the mechanisms regulating their neurochemical and peptidergic output to GnRH neurons, and how these contribute to GnRH/LH surge generation.

## Abbreviations

LH: luteinizing hormone
GnRH: gonadotropin-releasing hormone
ERα: estrogen receptor alpha
AVPV: anteroventral periventricular region
4-AP: 4-aminopyridine
TEA: tetraethylammonium chloride
F-I: firing vs current
V_50_: membrane potential at half activation or inactivation
g_max_: maximum conductance

## Acknowledgements

We thank Elizabeth Wagenmaker and Laura Burger for expert technical assistance and editorial comments. We thank James L. Kenyon, University of Nevada, Reno, for the Excel spreadsheet used to calculate junction potentials.

